## Supplementary text, figures and tables for "A solution to the learning dilemma for recurrent networks of spiking neurons"

### S1 Eligibility traces

Eligibility traces have been introduced in section "Mathematical basis for *e-prop*" in Results. Here, we provide further information on eligibility traces. In section S1.1, we discuss an alternative view on eligibility traces as derivatives. Second, we extend in section S1.3 our treatment of eligibility traces for LSNNs in Methods to include non-uniform synaptic delays.

#### S1.1 Viewing eligibility traces as derivatives

The notion of derivative $\left[\frac{d\mathbf{h}_j^t}{dW_{ji}}\right]_{\text{local}}$ from equation (14) quantifies the influence of an infinitesimal change of $W_{ji}$ on the hidden state $\mathbf{h}_j^t$ through the internal processes of neuron $j$ and the spikes of neuron $i$. Unlike the partial derivative $\frac{\partial \mathbf{h}_j^t}{\partial W_{ji}}$ it does not only take into account the update of the hidden state at time step $t$, but considers instead the full neuron history. In comparison to the total derivative $\frac{d\mathbf{h}_j^t}{dW_{ji}}$ it is only aware of the activity of neurons $i$ and $j$. Below we provide a formal explanation why the eligibility traces and the eligibility vectors can be viewed as derivatives.

One arrives at the equation (13) if one computes the total derivative $\frac{d\boldsymbol{h}_j^t}{dW_{ji}}$ in a variation of the computational graph of Fig. 6 where the function $\mathbf{h}_j^t = M(\mathbf{h}_j^{t-1}, \widetilde{\mathbf{z}}^{t-1}, \mathbf{x}^t, \mathbf{W}_j)$ receives a vector $\widetilde{\mathbf{z}}^{t-1}$ that is equal to $\mathbf{z}^{t-1}$ but considered as a constant. We can then define the notation $\left[\frac{dA}{dB}\right]_{\text{local}}$ as a total derivative $\frac{dA}{dB}$ in this altered computational graph. Following the same derivation as for equation (15) in Methods, the total derivative $\left[\frac{d\mathbf{h}_j^t}{dW_{ji}}\right]_{\text{local}}$ expends as $\sum_{t\leq t'} \left[\frac{d\mathbf{h}_j^{t'}}{d\mathbf{h}_j^t}\right]_{\text{local}} \frac{\partial h_j^t}{\partial W_{ji}}$. Viewing that $\left[\frac{d\mathbf{h}_j^t}{d\mathbf{h}_j^{t-1}}\right]_{\text{local}} = \frac{\partial \mathbf{h}_j^t}{\partial \mathbf{h}_j^{t-1}}$ in this computational graph, one recognizes that $\left[\frac{d\mathbf{h}_j^t}{dW_{ji}}\right]_{\text{local}}$ is the eligibility vector given in equation (14). Equation (13) follows since $e_{ji}^t = \frac{\partial z_j^t}{\partial \mathbf{h}_j^t} \cdot \epsilon_{ji}^t = \left[\frac{dz_j^t}{dW_{ji}}\right]_{\text{local}}$. In fact this definition allows the extension of this new local derivative to other quantities and one can summarize *symmetric e-prop* as the replacement of $\frac{dE}{dW_{ji}}$ by $\left[\frac{dE}{dW_{ji}}\right]_{\text{local}}$ in stochastic gradient descent.

#### S1.2 Eligibility traces for LSNNs with membrane potential reset

The eligibility traces derived in the methods do not take the reset term into account. We derive here the eligibility traces that can correct for this. Note however that we did not observe an improvement when using this more complex model on the phoneme recognition and the task where temporal credit assignment is difficult.

**Eligibility traces for LIF neurons.** When taking into account the reset, the partial derivative $\frac{\partial \boldsymbol{h}_j^{t+1}}{\partial \boldsymbol{h}_j^t}$ becomes $\alpha - v_{\text{thr}}\psi_j^t$ instead of $\alpha$ and, accordingly to equation (14), the eligibility vector can be computed with the recursive formula: $\boldsymbol{\epsilon}_{ji}^{t+1} = (\alpha - \beta\psi_j^t)\boldsymbol{\epsilon}_{ji}^t + z_j^t$.

**Eligibility traces for ALIF neurons.** According to the dynamics of the ALIF neurons defined in equations (6)–(10) one coefficient differs in the matrix $\frac{\partial \boldsymbol{h}_j^{t+1}}{\partial \boldsymbol{h}_j^t} \in \mathbb{R}^{2\times 2}$ as soon as one takes the reset into account. The coefficient $\frac{\partial v_j^t}{\partial a_j^t}$ was 0 without reset and becomes now $v_{\text{thr}}\beta\psi_j^t$. Overall the full derivative $\frac{\partial \boldsymbol{h}_j^{t+1}}{\partial \boldsymbol{h}_j^t}$ is then equal to:

$$\frac{\partial \boldsymbol{h}_j^{t+1}}{\partial \boldsymbol{h}_j^t} = \begin{pmatrix} \alpha - v_{\text{thr}}\psi_j^t & v_{\text{thr}}\beta\psi_j^t \\ \psi_j^t & \rho - \beta\psi_j^t \end{pmatrix} . \tag{S1}$$

Even-though *e-prop* can be implemented as such, the recursive propagation of the eligibility vector in equation (14) cannot be written in the form of two separable equations as done in equations (24) and (25). Hence, we preferred to ignore the reset in Methods to provide more interpretable equations for eligibility traces.

##### S1.3 Eligibility traces for LSNNs with non-uniform synaptic delays

In our derivation of eligibility traces for LSNNs, we used uniform synaptic delays to ease notation. Here, we detail how *e-prop* can be extended to non-uniform delays of several ms. Let the delay of a synapse from neuron $i$ to $j$ be denoted by $c(j,i) > 0$. Similarly, let $d(j,i) \geq 0$ be the delay of a synapse that connects an input neuron $i$ with neuron $j$. Using this definition, the dynamics of the membrane potential, see equation (6), is written as:

$$v_j^{t+1} = \alpha v_j^t + \sum_{i \neq j} W_{ji}^{\text{rec}} z_i^{t-c(j,i)} + \sum_i W_{ji}^{\text{in}} x_i^{t-d(j,i)} - z_j^t v_{\text{th}} \ . \tag{S2}$$

Like in the uniform delay case, we obtain $\frac{\partial v_j^{t+1}}{\partial v_j^t} = \alpha$. The difference for arbitrary delays becomes visible in $\frac{\partial v_j^t}{\partial W_{ji}^{\text{rec}}} = z_i^{t-c(j,i)}$ and in $\frac{\partial v_j^t}{\partial W_{ji}^{\text{in}}} = x_i^{t-d(j,i)}$. For recurrent weights, the component of the eligibility vector associated to the membrane potential is hence:

$$\epsilon_{ji,v}^t = \sum_{t' \leq t-c(j,i)} z_i^{t'} = \bar{z}_i^{t-c(j,i)} \ . \tag{S3}$$

As the dynamics of the threshold adaptation is unchanged, the update of $\epsilon_{ji,a}^t$ remains as given in equation (24). We obtain an eligibility trace

$$e_{ji}^t = \psi_j^t \left( \bar{z}_i^{t-c(j,i)} - \beta \epsilon_{ji,a}^t \right) \ . \tag{S4}$$

Analogously, we obtain the corresponding eligibility trace for input synapses by replacing $z_i^t$ and $c(j,i)$ with $x_i^t$ and $d(j,i)$ respectively.

#### S2 Optimization and regularization procedures

Here, we discuss how optimization of networks was implemented and techniques that were used to regularize networks.

##### S2.1 Optimization procedure

For *e-prop* and for *BPTT*, the weights were updated once after a batch of training trials. For simplicity, all the weight updates $\Delta W_{ji}^{\text{rec}}$ are written for the most basic version of stochastic gradient descent ($\Delta W_{ji}^{\text{rec}} = -\eta \widehat{\frac{dE}{dW_{ji}^{\text{rec}}}}$, where $\widehat{\frac{dE}{dW_{ji}^{\text{rec}}}}$ is the gradient estimate) in this article. In practice, we used Adam [2] to boost stochastic gradient descent. We refer to [2] for the computation of the weight updates that result from the gradient estimates.

##### S2.2 Firing rate regularization for LSNNs

To ensure a low firing rate in LSNNs, we added a regularization term $E_{\text{reg}}$ to the loss function $E$. This regularization term had the form:

$$E_{\text{reg}} = \frac{1}{2} \sum_j \left( f_j^{\text{av},t} - f^{\text{target}} \right)^2 \ , \tag{S5}$$

where $f^{\text{target}}$ is a target firing rate and $f_j^{\text{av}} = \frac{1}{Tn_{\text{trial}}} \sum_t z_j^t$ is the average firing rate of neuron $j$. Here, the sum runs over the time steps of all the $n_{\text{trial}}$ trials between two weight updates. To derive the plasticity rule that implements this regularization, we follow equation (27) in Methods. The partial derivative of the regularization loss has the form:

$$\frac{\partial E_{\text{reg}}}{\partial z_j^t} = \frac{1}{Tn_{\text{trial}}} \left( f_j^{\text{av}} - f^{\text{target}} \right) . \tag{S6}$$

Inserting this expression into equation (27), we obtain the plasticity rule that implements the regularization:

$$\Delta W_{ji}^{\text{rec}} \quad = \eta \, c_{\text{reg}} \textstyle\sum_t \frac{1}{Tn_{\text{trial}}} \left( f^{\text{target}} - f_j^{\text{av}} \right) e_{ji}^t \ , \tag{S7}$$

where $c_{\text{reg}}$ is a positive coefficient that controls the strength of the regularization. This plasticity rule is applied simultaneously together with the plasticity rule that minimizes the loss $E$. Note that this weight update fits the *e-prop* framework provided by equation (1) with a learning signal $L_j^{\text{reg}}$ proportional to $f^{\text{target}} - f_j^{\text{av}}$ available locally at neuron $j$. The learning signal $L_j^{\text{reg}}$ can be computed locally and added to the task-specific learning signal $L_j^t$. A variant of this regularization replaces the average firing rate $f_j^{\text{av}}$ with a running average $f_j^{\text{av},t}$ that accumulates the spikes since the last weight update. It can achieve the same effect with less resources when implemented online.

#### S2.3 *Adaptive e-prop* and weight decay regularization

In *adaptive e-prop* all neurons are receiving learning signals with random broadcast weights $B_{jk}$ as in *random e-prop*. Yet, if neuron $j$ sends activity to the output neuron $k$ via output weights $W_{kj}^{\text{out}}$, the broadcast weights $B_{jk}$ are subsequently adapted. This was implemented by applying an identical weight update to the broadcast and the output weights, i.e. $\Delta B_{jk} = \Delta W_{kj}^{\text{out}}$ and by subtracting $c_{\text{decay}} \cdot W_{kj}^{\text{out}}$ (resp. $c_{\text{decay}} \cdot B_{jk}$) from the weight $W_{kj}^{\text{out}}$ (resp. $B_{jk}$) after each weight update, where $c_{\text{decay}} > 0$ is the regularization factor (see specific experiments for the value of $c_{\text{decay}}$). This weight decay in combination with the mirroring of the weight updates has the effect that, despite different initialization, the output weights and the adaptive broadcast weights converge to similar values. The remaining difference of performance between *symmetric* and *adaptive e-prop* reported in Fig. S2 and Fig. S4 may be explained by the different initializations.

*Adaptive e-prop* can be viewed as that version of *e-prop* that exploits all types of biologically plausible learning signals that are available at individual neurons, apart from reward prediction errors or signals from a separately trained RSNN that sends learning signals, see Fig. 3 of [3].

#### S2.4 Voltage regularization

One advantage of a gradient-based learning method is that additional terms can be added to the loss function in order to enforce a particular network behavior. We utilized this possibility to regularize the membrane voltage of network neurons, using an idea of Arjun Rao. This regularization reduced excessive membrane voltages (which can otherwise even exceed the threshold voltage during the neuron's refractory period) and led to sparser and more irregular firing activity, similar as in biological data. In particular, we used a regularization term that penalized membrane potentials higher than the threshold voltage $A_t$ for ALIF neurons (resp. $v_{\text{thr}}$ for LIF) or lower than $-A_t$ for ALIF neurons (resp. $-v_{\text{thr}}$ for LIF). We add a regularization loss of the form:

$$E_{V,\text{reg}} = E_{\text{reg},V+} + E_{\text{reg},V-} = \frac{1}{2} \sum_t \left( \left[ v_j^t - A_j^t \right]^+ \right)^2 + \left( \left[ -v_j^t - A_j^t \right]^+ \right)^2 \tag{S8}$$

for ALIF neurons (replace $A_t$ by $v_{\text{thr}}$ for LIF neurons), where $[\cdot]^+$ is the rectified linear function. Unlike other loss functions, $E_{V,\text{reg}}$ depends directly on the voltage and not on the spikes. Hence instead of the full *e-prop* formula we use that the cost function is already local to neuron $j$, so that we can derive the weight update as follows (we consider only the first part $E_{\mathrm{reg},V+}$ of the loss function to simply the analysis):

$$\Delta W_{ji} = -\eta c_{\mathrm{reg},V} \sum_t \left[\frac{dE_{\mathrm{reg},V+}}{dW_{ji}}\right]_{\mathrm{local}} \tag{S9}$$

$$= -\eta c_{\mathrm{reg},V} \sum_t \frac{\partial E_{\mathrm{reg},V+}}{\partial \boldsymbol{h}_j^t} \boldsymbol{\epsilon}_j^t \tag{S10}$$

$$= -\eta c_{\mathrm{reg},V} \sum_t \left[v_j^t - A_j^t\right]^+ \left(\epsilon_{ji,v}^t - \epsilon_{ji,a}^t\right) \tag{S11}$$

$$= -\eta c_{\mathrm{reg},V} \sum_t \left[v_j^t - A_j^t\right]^+ \left(\bar{z}_i^{t-1} - \epsilon_{ji,a}^t\right) . \tag{S12}$$

##### S2.5 Optimization with rewiring for sparse network connectivity

Due to limited resources, neural networks in the brain and in neuromorphic harware are sparsely connected. In addition, the connectivity structure of brain networks is dynamic, with synaptic connections being added and deleted on the time scale of hours or days, which was shown to help the network to use the limited connectivity resources in an optimal manner [4]. In order to test whether *e-prop* is compatible with synaptic rewiring, we combined it with DEEP R [5]. DEEP R is based on a model for synaptic rewiring in the brain [4] and allows to rewire sparse neural network models during training with gradients descent. The algorithm minimizes the loss function $E$ subject to a constraint on the total number of connected synapses. To do so, each synaptic weight $W_{ji}$ is assigned a fixed sign $s_{ji}$ (it is defined to be excitatory or inhibitory) and an amplitude $w_{ji}$. Each potential synaptic connection can either be "active", i.e., the synaptic connection is realized, or "dormant", i.e., this potential connection is not realized.

For a dormant synaptic connection, the weight $W_{ji}$ is set to be zero and the gradients and weight updates of the connection $i \to j$ are not computed. It means in *e-prop* that dormant synapses do not require eligibility traces. For an active connection, the weight is defined as $W_{ji} = s_{ji} w_{ji}$ and the weight amplitude is updated according to the update $\Delta w_{ji} = s_{ji} \Delta W_{ji} - \eta c_{L1}$ where $\Delta W_{ji}$ is the weight update given here by *e-prop* and $c_{L1} = 0.01$ is an $L1$ regularization coefficient. To update the network structure such that the set of active connections is optimized along side their synaptic weights, DEEP R proceeds as follows after each weight update:

- every active connection for which the amplitude becomes negative is set to be dormant,
- and some dormant connections are selected randomly and set to be active with $w_{ji} = 0$ such that the total number of active connection remains constant.

We define the synapse signs $s_{ji}$ such that 80% of the neurons are excitatory and 20% are inhibitory. Despite the constraint on the neuron signs and the constraint that 90% of the synapses should remain dormant throughout the learning process, *e-prop* and rewiring solve the task where temporal credit assignment is difficult of Fig. 3.

#### S3 Supervised learning with *e-prop*

##### S3.1 Synaptic plasticity rules for *e-prop* in supervised learning

Here, we derive synaptic plasticity rules that result from *e-prop* for supervised learning. We consider two cases: First, we derive plasticity rules for regression tasks, and second, for classification tasks.

We follow the scheme described by equation (27) in Methods. Hence the loss gradients $\frac{dE}{dW_{ji}}$ are estimated using the approximation $\widehat{\frac{dE}{dW_{ji}}} \stackrel{\mathrm{def}}{=} \sum_t \frac{\partial E}{\partial z_j^t} e_{ji}^t$. Given the eligiblity traces that are derived in Methods and section S4.4, what remains to be derived for each task is the expression of the relevant derivative $\frac{\partial E}{\partial z_j^t}$ and show that it can be computed online.

**Regression tasks:** Consider a regression problem with loss function $E = \frac{1}{2}\sum_{t,k}(y_k^t - y_k^{*,t})^2$, targets $y_k^{*,t}$ and outputs $y_k^t$ as defined in equation (11). The partial derivative $\frac{\partial E}{\partial z_j^t}$ takes the form:

$$E = \frac{1}{2}\sum_{t,k}(y_k^t - y_k^{*,t})^2 \tag{S13}$$

$$\frac{\partial E}{\partial z_j^t} = \sum_k W_{kj}^{\text{out}} \sum_{t' \geq t}(y_k^{t'} - y_k^{*,t'})\kappa^{t'-t} \ . \tag{S14}$$

This seemingly provides an obstacle for online learning, because the partial derivative is a weighted sum over future errors. But this problem can be resolved. Following equation (1), the approximation $\widehat{\frac{dE}{dW_{ji}}}$ of the loss gradient is computed with *e-prop* as follows (we insert $\frac{\partial E}{\partial z_j^t}$ in place of the total derivative $\frac{dE}{dz_j^t}$):

$$\widehat{\frac{dE}{dW_{ji}}} = \sum_{t'} \frac{\partial E}{\partial z_j^{t'}} e_{ji}^{t'} \tag{S15}$$

$$= \sum_{k,t'} W_{kj}^{\text{out}} \sum_{t \geq t'}(y_k^t - y_k^{*,t})\kappa^{t-t'} e_{ji}^{t'} \tag{S16}$$

$$= \sum_{k,t} W_{kj}^{\text{out}} (y_k^t - y_k^{*,t}) \underbrace{\sum_{t' \leq t} \kappa^{t-t'} e_{ji}^{t'}}_{\stackrel{\text{def}}{=} \bar{e}_{ji}^t} \ , \tag{S17}$$

where we changed the order of summations in the last line. The second sum indexed by $t'$ is now over previous events that can be computed online. It is just a low-pass filtered version of the eligibility trace $e_{ji}^t$. With this additional filtering of the eligibility trace with a time constant equal to that of the leak of output neurons, we see that *e-prop* takes into account the latency between an event at time $t'$ and its impact on later errors at time $t$ within the integration time window of the output neuron. Hence, implementing weight updates with gradient descent and learning rate $\eta$, the plasticity rule resulting from *e-prop* is given by the equation (28). The gradient of the loss function with respect to the output weights $\frac{dE}{dW_{kj}^{\text{out}}}$ can be implemented online without relying on the theory of *e-prop*. The plasticity rule resulting from gradient descent is directly:

$$\Delta W_{kj}^{\text{out}} = -\eta \sum_t (y_k^t - y_k^{*,t}) \mathcal{F}_\kappa(z_j^t) \ . \tag{S18}$$

Similarly the update of the bias of the output neurons is $\Delta b_k^{\text{out}} = -\eta \sum_t (y_k^t - y_k^{*,t})$.

**Classification tasks:** We assume that $K$ target categories are provided in the form of a $K$-dimensional one-hot encoded vector $\boldsymbol{\pi}^{*,t}$. To train recurrent networks in this setup, we replace the mean squared error by the cross entropy loss:

$$E = -\sum_{t,k} \pi_k^{*,t} \log \pi_k^t \ , \tag{S19}$$

where the probability for class $k$ predicted by the network is given as $\pi_k^t = \text{softmax}_k(y_1^t, \dots, y_K^t) = \exp(y_k^t)/\sum_{k'}\exp(y_{k'}^t)$. To derive the modified learning rule that results from this loss function $E$, we replace $\frac{\partial E}{\partial z_j^t}$ of equation (S14) with the corresponding one resulting from (S19):

$$\frac{\partial E}{\partial z_j^t} = \sum_k W_{kj}^{\text{out}} \sum_{t' \geq t}(\pi_k^{t'} - \pi_k^{*,t'})\kappa^{t'-t}. \tag{S20}$$

Following otherwise the same derivation as in equations (S15)-(S17), the plasticity rule in the case of classification tasks is given by equation (29).

Similarly, one obtains the plasticity rule for the output connections, where the only difference between the cases of regression and of classification is that the output $y_k^t$ and the target $y_k^{*,t}$ are replaced by $\pi_k^t$ and $\pi_k^{*,t}$ respectively: $\Delta W_{kj}^{\text{out}} = -\eta \sum_t (\pi_k^t - \pi_k^{*,t}) \mathcal{F}_\kappa(z_j^t)$. The update of the bias of the output neurons is $\Delta b_k^{\text{out}} = -\eta \sum_t (\pi_k^t - \pi_k^{*,t})$.

#### S3.2 Simulation details: phoneme recognition task (Fig. 2)

##### S3.2.1 Frame-wise phoneme classification

The goal of the frame-wise setup of the task is to classify audio-frames into phoneme classes. Every input sequence of audio-frames has a corresponding sequence of class labels of the same length, hence the model does not need to align the input sequence to the target sequence. This task has been widely adopted as a phoneme recognition benchmark for recurrent neural networks (RNNs).

**Details of the network model:** In Fig. 2c the performance is reported for the same network architecture as used for LSTM networks in [6], it uses in particular a bi-diretional architecture where the output of the LSNN is augmented by the output of a second LSNN that receives the input sequence in reverse time order. This reduces the error in comparison to the uni-directional case from 36.1% to 32.9% for LSNNs with *BPTT*. This improvement is qualitatively similar to what was previously reported for LSTM networks [6].

With the bi-directional architecture used in Fig. 2c, each of the two networks consisted of 300 LIF neurons and 100 ALIF neurons. The neurons in the LSNN had a membrane time constant of $\tau_m = 20$ ms, an adaptation time constant of $\tau_a = 200$ ms, an adaptation strength of $\beta = 0.184$, a baseline threshold $v_{\text{th}} = 1.6$, and a refractory period of 2 ms.

We used 61 output neurons in total, one for each class of the TIMIT dataset. The membrane time constant of the output neurons was $\tau_{out} = 3$ ms. A softmax was applied to their output, resulting in the corresponding class probabilities. The network model had $\approx 0.4$ million weights.

**Details of the dataset preparation and of the input preprocessing:** We followed the same task setup as in [7, 6]. The TIMIT dataset was split according to [8] into a training, validation, and test set with 3696, 400, and 192 sequences respectively. The input $\boldsymbol{x}^t$ was given as preprocessed audio that was obtained by the following procedure: Computation of 13 Mel Frequency Cepstral Coefficients (MFCCs) with a frame size of 10 ms on an input window of length 25 ms, computation of the first and the second derivatives of MFCCs, concatenation of all computed factors. The 39 input channels were mapped to the range $[0, 1]$ according to the minimum/maximum values in the training set.

In order to map the inputs into the temporal time domain of LSNNs, each preprocessed audio frame was fed as inputs $\boldsymbol{x}^t$ to the LSNN for 5 consecutive 1 ms steps.

**Details of the learning procedure:** All networks were trained for a maximum of 80 epochs, where we used early stopping to report the test error at the point of the lowest error on the validation set. Weight updates were implemented using Adam with default hyperparameters [2] except for $\epsilon_{\text{Adam}}$, which was set to $10^{-5}$. Gradients were computed using batches of size 32. We used L2 regularization in all networks by adding the term $10^{-5} \cdot \|W\|^2$ to the loss function, where $W$ denotes all weights in the network. The learning rate was initialized to 0.01 and fixed during training. For *random e-prop* and *adaptive e-prop*, broadcast weights $B_{jk}$ were initialized using a Gaussian distribution with a mean of 0 and a variance of 1 and $1/n$ ($n$ being the number of neurons in the LSNN) respectively. In *adaptive e-prop*, we used in addition to the weight decay described above L2 weight decay on output and broadcast weights according to S2.3 using a factor of $c_{\text{decay}} = 10^{-2}$. Firing rate regularization, as described in section S2.2, was applied with $c_{\text{reg}} = 50$.

##### S3.2.2 Phoneme sequence recognition with CTC

We compared *e-prop* and *BPTT* on the task and the network architecture used in [9]. The essential building blocks of this architecture were also used in [10] for developing commercial software for speech-to-text transcriptions. In this architecture Connectionist Temporal Classification (CTC) is employed. This enabled us to train networks on unaligned sequence labeling tasks end-to-end. We considered the results of [9] that were obtained with three layers of bi-directional LSTM networks, CTC, and *BPTT* as a reference. We are aware that this configuration cannot be adapted to an online implementation easily, due to the usage of a bi-directional LSTM network and the CTC loss function. However, we believe that this task is still relevant to compare *BPTT* and *e-prop* because it is a well established benchmark for RNNs.

**Details of the network model:** The neurons were structured into 3 layers. The network was recurrently connected within a layer and had feedforward connections across layers. Each layer consisted of 80 LIF neurons and 720 ALIF neurons (9.1 million weights). The neurons in LSNNs had a membrane time constant of $\tau_m = 20$ ms, an adaptation time constant of $\tau_a = 500$ ms, an adaptation strength of $\beta = 0.074$, a baseline threshold $v_{\text{th}} = 0.2$, and a refractory period of 2 ms. Synaptic delays were randomly chosen from $\{1, 2\}$ ms with equal probability. The membrane time constant of output neurons was $\tau_{out} = 3$ ms.

***E-prop* with many layers of recurrent neurons:** If one naively applies *e-prop* in such a configuration, the partial derivative $\frac{\partial E}{\partial z_j^t}$ is non-zero only if $j$ belongs to the last layer, whereas earlier layers would not receive any learning signal. To avoid this, we connected all neurons in all layers of the RNN to the output neurons. Therefore, the outputs $y_k^t$ of the RNN were given as $y_k^t = \sum_{t' \leq t} \kappa^{t-t'} \sum_l \sum_j W_{kj}^{\text{out},(l)} z_j^{(l),t'}$, where $z_j^{(l),t'}$ denotes the visible state of a neuron $j$ within the layer $l$. As a result, the learning signals in the case of *e-prop* were non-zero for neurons in every layer.

***E-prop* with the CTC loss function:** $E_{CTC}$ is defined based on the log-likelihood of obtaining the sequence of labeled phonemes given the network outputs $y_k^t$. We refer to [11] for the formal definition of the probabilistic model. Equation (7.27) in [12] shows the gradient of the loss function $E_{CTC}$ with respect to the activity of the outputs $y_k^t$ that we denote as $\frac{dE}{dy_k^t}$. Using the linear relationship between the visible state $z_j^{(l),t}$ and the outputs $y_k^t$, we obtain that the partial derivative $\frac{\partial E_{CTC}}{\partial z_j^{(l),t}}$ that we need in order to find the learning signals used in *e-prop*, and are defined as $\sum_{t' \geq t} \kappa^{t'-t} \sum_k \frac{dE}{dy_k^{t'}} B_{jk}^{(l)}$. Here, $B_{jk}^{(l)}$ denote the broadcast weights to the layer $l$.

**Details of the dataset preparation and of the input preprocessing:** The TIMIT dataset was split in the same manner as in the frame-wise version of the task. The raw audio was preprocessed before it was provided as an input $\boldsymbol{x}^t$ to the network. This included the following steps: computation of a Fourier-transform based filter-bank with 40 coefficients and an additional channel for the signal energy (with step size 10 ms and window size 25 ms), computation of the first and the second derivatives, concatenation of all computed factors, which totals to 123 input channels. Normalization over the training set was done in the same manner as in the frame-wise version of the task.

In order to map the inputs into the temporal time domain of LSNNs, each preprocessed audio frame was fed as inputs $\boldsymbol{x}^t$ to the LSNN for 5 consecutive 1 ms steps.

**Details of the learning procedure:** All models were trained for a total of 60 epochs, where gradients were computed using batches of 8 sequences. The learning rate was initialized to $10^{-3}$ and decayed every 15 epochs by a factor of 0.3. We used early stopping to report the test error, as in the previous task. Dropout was applied during training between the hidden layers and at the output neurons with a dropout probability of 0.3. As in the frame-wise setup, the weight updates were implemented using Adam with the default hyperparameters [2] except for $\epsilon_{\mathrm{Adam}} = 10^{-5}$. For *random e-prop* and *adaptive e-prop*, broadcast weights $B_{jk}$ were initialized using a Gaussian distribution with a mean of 0 and a variance of 1 and $1/n$ ($n$ denoting the number of neurons in the corresponding layer) respectively. In *adaptive e-prop*, we used L2 weight decay on output and broadcast weights according to S2.3 using a factor of $c_{\mathrm{decay}} = 10^{-4}$. When the global norm of gradients $N_{\mathrm{clip}} = \|\widehat{\frac{dE}{dW^{\mathrm{in}}_{ji}}}\|^2 + \|\widehat{\frac{dE}{dW^{\mathrm{rec}}_{ji}}}\|^2 + \|\widehat{\frac{dE}{dW^{\mathrm{out}}_{ji}}}\|^2$ was larger then 1, we scaled the gradients by a factor of $\frac{1}{N_{\mathrm{clip}}}$. We used beam search decoding with a beam width of 100. As in [9], the networks were trained on all 61 phoneme labels but were then mapped to a reduced phoneme set (39 classes) for testing.

#### S3.3 Applying *e-prop* to an episodic memory task

The FORCE training method [1] arguably defines the state of the art for training methods for RSNNs that do not need to backpropagate gradients through time. FORCE learning uses a synaptic plasticity rule that requires knowledge of the values of all synaptic weights in the network. This rule was not argued to be biologically plausible, but no other methods for training an RSNN to solve the task described below were known so far.

In order to compare *e-prop* to FORCE learning, we tested *e-prop* on the task to replay a movie segment that was used as a target $\boldsymbol{y}^{*,t}$ [1]. Specifically, it had to generate at each time step the values of all pixels that described the video frame of the movie at that time step. This episodic memory task was arguably the most difficult task for which an RSNN was previously trained in [1].

Here, we considered an extension to this task: An LSNN had to replay 1 out of 3 possible movies, where the desired movie index was provided as a cue to the network, see Fig. S1a. As in [1], the LSNN received also a clock-like input signal to indicate the current position in the movie. We show in Fig. S1b that an LSNN can be trained to solve this task with either one of the *e-prop* versions (see Movie S4), and that *e-prop* performs almost as well as *BPTT*.

**Details of the network model:** We used an LSNN that consisted of 700 LIF neurons and 300 ALIF neurons. Each neuron had a membrane time constant of $\tau_m = 20$ ms and a refractory period of 5 ms. ALIF neurons had a threshold adaptation time constant of 500 ms, and a threshold adaptation strength of $\beta = 0.07$. All neurons had a baseline threshold of $v_{\mathrm{th}} = 0.62$. All 5544 output neurons had a membrane time constant of $\tau_{out} = 4$ ms.

**Details of the dataset preparation and of the input scheme:** We manually chose three movie clips from the Hollywood 2 dataset [13], which contained between 0 and 2 scene cuts*, see Movie S4. The movie clips were clipped to a length of 5 seconds and spatially subsampled to a resolution of $66 \times 28$ pixels. Since our simulations used 1 ms as a discrete time step, we linearly interpolated between the frames of the original movie clips, which had a framerate of 25 frames per second. In total, we obtained a target signal with $66 \times 26 \times 3 = 5544$ dimensions, whose values were divided by a constant of 255, such that they fit in the range of $[0, 1]$.

The network received input from 115 input neurons, divided into 23 groups of 5 neurons. The first 20 groups indicated the current phase of the target sequence, similar to [1]. Neurons in group $i \in \{0, 19\}$ produced regular spike trains with a firing rate of 50 Hz during the time interval $[250 \cdot i, 250 \cdot i + 250)$ ms and were silent at other times. The remaining 3 groups encoded which movie had to be replayed, where each group was assigned to one of the three movies. To indicate a desired replay of one specific movie, each neuron in the corresponding group produced a Poisson spike train with a rate of 50 Hz and was silent otherwise.

*sceneclipautoautotrain00019.avi, sceneclipautoautotrain00061.avi, sceneclipautoautotrain00071.avi

**Details of the learning procedure:** For learning, we carried out 5 second simulations, where the network produced a 5544 dimensional output pattern. Gradients were accumulated for 8 successive trials, after which weight updates were applied using Adam with a learning rate of $2 \cdot 10^{-3}$ and default hyperparameters [2]. The movie to be replayed in each trial was selected with uniform probability. After every 100 weight updates (iterations), the learning rate was decayed by a factor of 0.95. For *random e-prop*, we used random broadcast weights $B_{jk}$ that were sampled from a Gaussian distribution with a mean of 0 and a variance of 1. In *adaptive e-prop* we used L2 weight decay (see section S2.3) for the broadcast weights $B_{jk}$ and the output weights $W_{ji}^{\text{out}}$ with a factor of $c_{\text{decay}} = 0.001$. To avoid an excessively high firing rate, regularization, as described in section S2.2, was applied with $c_{\text{reg}} = 0.1$ and a target firing rate of $f^{\text{target}} = 10$ Hz.

#### S3.4 Simulation details: task where temporal credit assignment is difficult (Fig. 3)

This task was inspired by the task performed by rodents in [14]. Each trial was split into three periods: the cue period, the delay period, and the decision period. During the cue period, the subject was stimulated with 7 successive binary cues ("left" or "right"), and had to take a corresponding binary decision ("left" or "right") during the decision period. The trial was considered a success if the decision matched the side that was most often indicated by the 7 cues. No action was required during the delay period. Each cue lasted for 100 ms and the cues were separated by 50 ms. The duration of the delay was distributed uniformly between 500 ms and 1500 ms, and the decision period lasted for 150 ms. The results from Fig. 3 were averaged over 20 different runs, where each run was characterized by a different random seed. For each run this resulted in different initialization of the weight matrices and broadcast weights as well as different realizations of random inputs.

**Details of the network model and input scheme:** We used an LSNN that consisted of 50 LIF neurons and 50 ALIF neurons. All neurons had a membrane time constant of $\tau_m = 20$ ms, a baseline threshold of $v_{\text{th}} = 0.6$, and a refractory period of 5 ms. The time constant of the threshold adaptation was set to $\tau_a = 2000$ ms, and its impact on the threshold was given as $\beta = 1.74 \cdot 10^{-2}$.

Input to this network was provided by 4 populations of 10 neurons each. The first two input populations encoded the cues as follows: When a cue indicated the "left" side (resp. the "right" side), all the neurons within the first (resp. the second) population produced Poisson spike trains with a firing rate of 40 Hz. The third input population spiked randomly throughout the decision period with a firing rate of 40 Hz and was silent otherwise. All the neurons in the last input population produced Poisson spike trains of 10 Hz throughout the trial, which was useful in particular to avoid that the network becomes quiescent during the delay.

**Details of the learning procedure:** For learning, we used *e-prop* for classification tasks, see section S3.1. The target label $\pi_k^{*,t}$ was given as the correct output during the decision period at the end of a trial. To help the network solve the task, we used a curriculum with an increasing number of cues. We first trained with a single cue, and increased the number of cues to 3, 5 and finally 7. The number of cues increased each time the network achieved less than 8% error on 512 validation trials. The same criterion was used to stop training once 7 cues were reached.

Independent of the learning algorithm that was used (*BPTT*, *e-prop*), a weight update was applied once every 64 trials and the gradients were accumulated during those trials additively. All weight updates were implemented using Adam with default parameters [2] and a learning rate of $5 \cdot 10^{-3}$. In the cases of *random e-prop* and *adaptive e-prop*, broadcast weights $B_{jk}$ were initialized using a Gaussian distribution with mean 0 and variance 1. In *adaptive e-prop* we used L2 weight decay (see section S2.3) for the broadcast weights $B_{jk}$ and the output weights $W_{ji}^{\text{out}}$ with a factor of $c_{\text{decay}} = 0.001$. In addition, firing rate regularization, as described in section S2.2, was applied with $c_{\text{reg}} = 1.$ and a target firing rate of $f^{\text{target}} = 10$ Hz.

**Details of the control experiment in Fig. S8:** For this control experiment, we introduced 1 to 3 additional feedforward (recurrent connections disabled) layers of 50 LIF neurons each before the output neurons. Each layer was connected in an all-to-all fashion to the preceding layer. All neurons had a membrane time constant of $\tau_m = 20$ ms. There was no voltage or firing rate regularization applied to the feedforward layers. We also applied the same curriculum of an increasing number of cues as described in the preceding section. In addition each stage of the curriculum was restricted to 1000 weight updates. All other hyperparameters were identical to the preceding section.

**Analysis of information in adaptive firing thresholds:** We verified (using 64 runs with 128 trials each for testing) that the firing threshold during the delay (500 ms after the last input cue) contained a substantial part of the decision-relevant information. The correct decision could be decoded by a simple readout network (1 hidden layer consisting of 100 ReLU neurons, trained over 200 iterations with 128 trials each) from the current values of the firing thresholds of ALIF neurons with an accuracy of $95.6\% \pm 1.82\%$ (standard deviation). From low pass filtered spikes of LIF neurons at the end of the recall period it could be decoded with an accuracy of $86.6\% \pm 3.7\%$ (standard deviation).

### S4 Applying supervised learning with *e-prop* to artificial neural networks (LSTM networks)

Here we show that *e-prop* can also be applied to artificial neural networks. We chose long short-term memory (LSTM) neworks [15] for this demonstration, whose performance defines the standard for RNNs in machine learning. We demonstrate in section S4.1 that LSTM networks can achieve competitive results on TIMIT when trained with *e-prop*, followed by details on these simulations (section S4.2). In the following sections, we provide details on the LSTM network model used (section S4.3) and on eligibility traces for LSTM units (section S4.4).

#### S4.1 Phoneme recognition with LSTM networks and *e-prop*

In Results, we have used *e-prop* to train LSNNs on the phoneme recognition task TIMIT (see Fig. 2). To test whether *e-prop* is effective also for artificial neural networks, we applied it to LSTM networks on the very same task in its two flavors of frame-wise classification and sequence transcription.

Supplementary Fig. S4 shows that *e-prop* approximates the performance of *BPTT* in both versions of TIMIT also for LSTM networks very well. For the more difficult version of TIMIT involving sequence transcription, we trained a feedforward sequence of 3 LSTM networks as in [9].

#### S4.2 Simulation details: phoneme recognition task with LSTM networks (Fig. S4)

The data preparation in the two setups (frame-wise phoneme classification and phoneme sequence recognition) were identical to the LSNN case. They are described in section S3.2. The details on the network models and training procedures are described next for the two task setups separately.

##### S4.2.1 Frame-wise phoneme classification with LSTM networks

**Details of the network model:** We used a bi-directional network architecture [6], where the output of an LSTM network was augmented by the output a second LSTM network that received the input sequence in reverse time order. Each of the two networks consisted of 200 LSTM units.

We used a 61-fold softmax output, one for each class of the TIMIT dataset. The LSTM network had $\approx 0.4$ million weights, which matched the number of weights in the LSNN for the same task.

**Details of the learning procedure:** LSTM networks were trained in the same way as LSNNs, see section S3.2, except for the following differences in hyperparameters: We decayed the learning rate after every 500 weight updates by a factor of 0.3. For L2 weight decay on output and broadcast weights according to S2.3 we used a factor of $c_{\text{decay}} = 10^{-3}$. As LSTM units are not spiking, we did not use firing rate regularization.

##### S4.2.2 Phoneme sequence recognition with CTC and LSTM networks

We compared *e-prop* and *BPTT* on the task and the network architecture used in [9]. As for LSNNs, we employed Connectionist Temporal Classification (CTC) to achieve phoneme sequence recognition (see "Phoneme sequence recognition with CTC" in section S3.2). This enabled us to train networks on unaligned sequence labeling tasks end to end.

**Details of the network model:** The neurons were structured into 3 recurrent layers. In each layer there were 250 LSTM units. All neurons in all layers of the RNN were connected to the output layer (see "*E-prop* with many layers of recurrent neurons" in section S3.2).

**Details of the learning procedure:** LSTM networks were trained in the same way as LSNNs, see section S3.2. In the case of *BPTT*, we also used the peephole feature in the LSTM network model.

#### S4.3 LSTM network model

We used a standard model for LSTM units [15], for which the hidden state at time step $t$ is a one dimensional vector containing only the content of the memory cell $c_j^t$, such that $\mathbf{h}_j^t \stackrel{\text{def}}{=} [c_j^t]$, and $z_j^t$ is the value of its output. The memory cell can be viewed as a register which supports writing, updating, deleting and reading. These operations are controlled independently for each cell $j$ at each time $t$ by input, forget and output gates (denoted by $i_j^t$, $f_j^t$ and $o_j^t$ respectively). The new cell state candidate that may replace the cell state $c_j^{t-1}$ at each time step $t$ is denoted $\widetilde{c}_j^t$. The input, forget, and output sigmoidal gates as well as the cell state candidate of an LSTM unit $j$ are defined by the following equations:

$$i_j^t = \sigma\big(\sum_i W_{ji}^{\text{rec},i} z_i^{t-1} + \sum_i W_{ji}^{\text{in},i} x_i^t\big) \quad \text{(S21)}$$

$$f_j^t = \sigma\big(\sum_i W_{ji}^{\text{rec},f} z_i^{t-1} + \sum_i W_{ji}^{\text{in},f} x_i^t\big) \quad \text{(S22)}$$

$$o_j^t = \sigma\big(\sum_i W_{ji}^{\text{rec},o} z_i^{t-1} + \sum_i W_{ji}^{\text{in},o} x_i^t\big) \quad \text{(S23)}$$

$$\widetilde{c}_j^t = \tanh\big(\sum_i W_{ji}^{\text{rec},c} z_i^{t-1} + \sum_i W_{ji}^{\text{in},c} x_i^t\big), \quad \text{(S24)}$$

where all the weights used here are parameters of the model (we also used biases that were omitted for readability). Using these notations, one can now write the update of the state of an LSTM unit $j$ in a form that we can relate to our general formalism:

$$c_j^t = f_j^t c_j^{t-1} + i_j^t \widetilde{c}_j^t \quad \text{(S25)}$$

$$z_j^t = o_j^t c_j^t \, . \quad \text{(S26)}$$

In terms of the computational graph in Fig. 6, equation (S25) defines $M(c_j^{t-1}, \mathbf{z}^{t-1}, \mathbf{x}^t, \mathbf{W}_j)$ and (S26) defines $f(c_j^t, \mathbf{z}^{t-1}, \mathbf{x}^t, \mathbf{W}_j)$.

#### S4.4 Eligibility traces for LSTM units

Eligibility traces for LIF neurons and ALIF neurons were derived in "Derivation of eligibility traces for concrete neuron models" in Methods. Here, we derive eligibility traces for the weights of LSTM units.

To obtain the eligibility traces, we note that the state dynamics of an LSTM unit is given by: $\frac{\partial \mathbf{h}_j^t}{\partial \mathbf{h}_j^{t-1}} = \frac{\partial c_j^t}{\partial c_j^{t-1}} = f_j^t$. For each weight $W_{ji}^{A,B}$ with $A$ being either "in" or "rec" and $B$ being $i, f$, or $c$, we compute a set of eligibility traces. For example, the eligibility vectors for the recurrent weights to the input gate $W_{ji}^{\mathrm{rec},i}$, are updated according to equation (14), leading to:

$$\boldsymbol{\epsilon}_{ji}^{i,t} = f_j^t \boldsymbol{\epsilon}_{ji}^{i,t-1} + \tilde{c}_j^t i_j^t (1 - i_j^t) z_i^{t-1} \ , \tag{S27}$$

resulting in eligibility traces:

$$e_{ji}^{i,t} = o_j^t \boldsymbol{\epsilon}_{ji}^{i,t} \ . \tag{S28}$$

Similarly, the eligibility traces for the input weights to the input gate are obtained by replacing $z_i^{t-1}$ with $x_i^t$.

**Output gates:** The gradients with respect to the parameters of the output gate do not require additional eligibility traces. This is because the output gate contributes to the observable state but not to hidden state, see equations S25 and S26. Therefore, one can use the standard factorization of the error gradient as used in *BPTT*. For the recurrent weights to the output gates $W_{ji}^{\mathrm{rec},o}$, the gradient is given by:

$$\frac{dE}{dW_{ji}^{\mathrm{rec},o}} = \sum_t \frac{dE}{dz_j^t} \frac{\partial z_j^t}{\partial W_{ji}^{\mathrm{rec},o}} = \sum_t \frac{dE}{dz_j^t} c_j^t o_j^t (1 - o_j^t) z_i^{t-1}. \tag{S29}$$

Hence, when applying *e-prop* to the weights of the output gate of an LSTM unit, we use, similarly as in equation (27), the same approximation $\frac{\partial E}{\partial z_j^t}$ of the ideal learning signal $\frac{dE}{dz_j^t}$. The other term is local, depending only on $t$ and $t-1$, and does not require eligibility traces. For input weights to the output gate $W_{ji}^{\mathrm{in},o}$, the gradient is obtained by replacing $z_i^{t-1}$ with $x_i^t$.

### S5 *Reward-based e-prop*: Application of *e-prop* to deep RL

#### S5.1 Synaptic plasticity rules for *reward-based e-prop*

Here, we derive the synaptic plasticity rules that result from gradients of the loss function $E$, as given in equation (32), see Fig. 4b for the network architecture. As a result of the general actor-critic framework with policy gradient, this loss function additively combines the loss function for the policy $E_\pi$ (actor) and the value function $E_V$ (critic).

We consider two cases: First, a simplified case where in each trial, one action is taken at the end of the trial. This is the setup of the reward-based version of the task where temporal credit assignment is difficult of Fig. 3, see Fig. S5 for performance results. Second, we analyse a more general case where actions are taken throughout the trial. This is the setup of the Atari tasks (Fig. 4, Fig. 5). For both cases, we derive the gradients for the parts $E_\pi$ and $E_V$ of the loss function $E$, and express the plasticity rules resulting from these gradients.

**Single action at the end of the trial (Fig. S5):** In this setup, a discrete action $a \in \{1, \ldots, K\}$ from a set of $K$ possibilities needs to be taken at the last time step $T$ of a trial, leading to a binary-valued reward $r^T$. As a result, the return $R^T$ (denoted here for notational simplicity as $R$) is equal to $r^T$. We assume that the agent chooses action $k$ with probability $\pi_k = \mathrm{softmax}_k(y_1^T, \ldots, y_K^T)$ $= \exp(y_k^T) / \sum_{k'} \exp(y_{k'}^T)$. Following the definition of $E_\pi$ in (30) one can write the loss function $E_\pi$ using the one-hot encoded action $\mathbb{1}_{a=k}$ which assumes a value of 1 only if $a = k$ and is 0 otherwise.

Hence we can write $\log \pi(a|\boldsymbol{y}) = \sum_k \mathbb{1}_{a=k} \log \pi_k$, where although we sum over all possible actions, only the term corresponding to the action $a$ that was taken is non zero. Therefore, we can write $E_\pi$ as:

$$E_\pi \quad = \quad -R \, \log \pi(a|\boldsymbol{y}) = -R \sum_k \mathbb{1}_{a=k} \log \pi_k \ . \tag{S30}$$

Interestingly, in the discrete action case, the loss function $E_\pi$ is reminiscent of the one used for supervised classification, see equation (S19). But it exhibits two differences: firstly, the indicator of the selected action $\mathbb{1}_{a=k}$ replaces the target label $\pi_k^*$, and secondly, the loss is multiplied by the reward $R$. Hence, the derivation of the weight update is almost identical with the additional difference that we also replace $R$ with $R - V$ in the resulting gradient to reduce the variance of the loss gradient estimate as derived in the general case in (35). Finally, in order to optimize $E$ defined in (32), we also need to consider $E_V = \frac{1}{2}(R - V)^2$, for which we can reuse the weight update of *e-prop* for regression (S17). Combining the weight updates for $E_\pi$ and $E_V$ into a single one, we have:

$$\Delta W_{ji}^{\text{rec}} \quad = \quad -\eta \underbrace{\left[ (R - V) \sum_k B_{jk}^\pi (\pi_k - \mathbb{1}_{a=k}) - c_V (R - V) B_j^V \right]}_{L_j} \, \bar{e}_{ji}^T \ , \tag{S31}$$

where we denote with $B_{jk}^\pi$ the broadcast weights from output neurons $y_k$, and with $B_j^V$ the broadcast weights from the output neuron that produces the value prediction $V$. The choice of these broadcast weights then defines which variant of *reward-based e-prop* is employed (*reward-based symmetric e-prop*, *reward-based adaptive e-prop*, or *reward-based random e-prop*).

For the synaptic connections of output neurons, the loss gradient can be computed directly from the loss function (32). We also subtract the value prediction to reduce variance of the gradient estimate as in (35), and obtain for the update rules: $\Delta W_{kj}^{\pi,\text{out}} = -\eta (R - V)(\pi_k - \mathbb{1}_{a=k}) \mathcal{F}_\kappa(z_j^T)$, and $\Delta W_j^V = \eta c_V (R - V) \mathcal{F}_\kappa(z_j^T)$. Similarly, the updates of the biases of output neurons are: $\Delta b_k^{\pi,\text{out}} = -\eta (R - V)(\pi_k - \mathbb{1}_{a=k})$, and $\Delta b^V = \eta c_V (R - V)$.

**Actions throughout the trial (Fig. 4, Fig. 5):** In this setup, we assume that the agent can take discrete actions $a_k$ among $K$ possible choices at certain decision times $t_0, \ldots, t_n, \ldots$. We also assume that the agent chooses action $k$ with a probability $\pi_k^{t_n}$ based on a categorical distribution that depends on the network output: $\pi_k^{t_n} = \pi(a_k|\boldsymbol{y}^{t_n}) = \text{softmax}_k(y_1^{t_n}, \ldots, y_K^{t_n}) = \exp(y_k^t) / \sum_{k'} \exp(y_{k'}^t)$.

To derive the gradient estimate resulting from *reward-based e-prop*, we first consider the regression problem defined by the loss function $E_V$, and note that a major difference to the previous case is that the return $R^t$ integrates future rewards that may arrive long after an action was taken. We begin with the result for regression from equation (S17). Substituting the relevant variables, we obtain an estimation of the loss gradient:

$$\widehat{\frac{dE_V}{dW_{ji}^{\text{rec}}}} = -\sum_{t'} (R^{t'} - V^{t'}) W_j^{V,\text{out}} \, \bar{e}_{ji}^{t'} \ , \tag{S32}$$

where $W_j^{V,\text{out}}$ are the weights of the output neuron $V_j^t$ predicting the value function $\mathbb{E}[R^t]$. In order to overcome the obstacle that an evaluation of the return $R^{t'}$ requires to know future rewards, we introduce temporal difference errors $\delta^t = r^t + \gamma V^{t+1} - V^t$, and use that $R^{t'} - V^{t'}$ is equal to the sum $\sum_{t \geq t'} \gamma^{t-t'} \delta^t$. We then reorganize the two sums over $t$ and $t'$ (note that the interchange of the summation order amounts to the equivalence between forward and backward view of RL [16]):

$$\widehat{\frac{dE_V}{dW_{ji}^{\mathrm{rec}}}} = -\sum_{t'}\Big(\sum_{t\geq t'}\gamma^{t-t'}\delta^t\Big)W_j^{V,\mathrm{out}}\,\bar{e}_{ji}^{t'} \tag{S33}$$

$$= -\sum_t \delta^t \sum_{t'\leq t}\gamma^{t-t'}W_j^{V,\mathrm{out}}\,\bar{e}_{ji}^{t'} \tag{S34}$$

$$= -\sum_t \delta^t\,\mathcal{F}_\gamma\big(W_j^{V,\mathrm{out}}\,\bar{e}_{ji}^{t}\big)\ . \tag{S35}$$

For the part $E_\pi$ in the loss function $E$ defined at equation (30), we consider the estimator $\widehat{\frac{\partial E}{\partial z_j^t}}$ given in (35), and use our previous definition that the probabilities of actions $a_k$ follow a categorical distribution, using a softmax of the network output $y_k$. The estimator then becomes:

$$\widehat{\frac{\partial E_\pi}{\partial z_j^t}} = \sum_k W_{kj}^{\pi,\mathrm{out}}\sum_{\{n\,|\,t_n\geq t\}}\kappa^{t_n-t}(R^{t_n}-V^{t_n})\left(\pi_k^{t_n}-\mathbb{1}_{a^{t_n}=k}\right)\ , \tag{S36}$$

where $W_{kj}^{\pi,\mathrm{out}}$ are the weights onto the output neurons $y_k^t$ defining the policy $\pi$, and $\kappa$ is the constant of the low-pass filtering of the output neurons. Following a derivation similar to equations (S15) to (S17) for supervised regression, we arrive at an estimation of the loss gradient of the form:

$$\widehat{\frac{dE_\pi}{dW_{ji}^{\mathrm{rec}}}} = \sum_t \widehat{\frac{\partial E_\pi}{\partial z_j^t}}e_{ji}^t \tag{S37}$$

$$= \sum_{t,k}W_{kj}^{\pi,\mathrm{out}}\sum_{\{n\,|\,t_n\geq t\}}(R^{t_n}-V^{t_n})\left(\pi_k^{t_n}-\mathbb{1}_{a^{t_n}=k}\right)\kappa^{t_n-t}e_{ji}^t \tag{S38}$$

$$= \sum_{n,k}(R^{t_n}-V^{t_n})W_{kj}^{\pi,\mathrm{out}}\left(\pi_k^{t_n}-\mathbb{1}_{a^{t_n}=k}\right)\underbrace{\sum_{t\leq t_n}\kappa^{t_n-t}e_{ji}^t}_{\bar{e}_{ji}^{t_n}}\ . \tag{S39}$$

Like in the derivation of the gradient of $E_V$, this formula hides a sum over future rewards in $R^{t_n}$ that cannot be computed online. It is resolved by introducing the backward view as in equation (S35). We arrive at the loss gradient:

$$\widehat{\frac{dE_\pi}{dW_{ji}^{\mathrm{rec}}}} = \sum_t \delta^t\mathcal{F}_\gamma\Big(\sum_k W_{kj}^{\pi,\mathrm{out}}\left(\pi_k^t-\mathbb{1}_{a^t=k}\right)\,\bar{e}_{ji}^t\Big)\ . \tag{S40}$$

Importantly, an action is only taken at times $t_0,\ldots,t_n,\ldots$, hence for all other times, we set the term $\pi_k^t-\mathbb{1}_{a^t=k}$ to zero.

Finally, the gradient of the loss function $E$ is the sum of the gradients of $E_\pi$ and $E_V$ in equations (S35) and (S40) respectively. Application of stochastic gradient descent with a learning rate of $\eta$ yields the synaptic plasticity rule given in the equations (36) and (37) in Methods.

**Policy entropy regularization:** In order to prevent premature convergence and encourage exploration, we add to the loss function of *reward-based e-prop* an auxiliary loss that maximizes the entropy $H$ of the policy by $E_H = -\sum_t H(\pi_1^t,\ldots,\pi_K^t)\overset{\mathrm{def}}{=}-\sum_t H^t$. Hence the equation (35) defining the estimators of the loss gradients $\widehat{\frac{\partial E}{\partial z_j^t}}$ becomes:

$$\widehat{\frac{\partial E}{\partial z_j^t}} = -\sum_n (R^{t_n}-V^{t_n})\frac{\partial\log\pi(a^{t_n}|\boldsymbol{y}^{t_n})}{\partial z_j^t}+c_V\frac{\partial E_V}{\partial z_j^t}+c_H\frac{\partial E_H}{\partial z_j^t}\ . \tag{S41}$$

To exhibit the term $\frac{\partial E_H}{\partial z_j^t}$, we begin with the definition of entropy and find $E_H = \sum_n \sum_k \pi_k^{t_n} \log \pi_k^{t_n} = -\sum_n H^{t_n}$ for the case of actions that probabilities distributed according to categorical distribution:

$$\frac{\partial E_H}{\partial z_j^t} = -\sum_{\{n \,|\, t_n \geq t\}} \sum_k \kappa^{t_n - t} W_{kj}^{\pi,\mathrm{out}} \frac{\partial H^{t_n}}{\partial y_k^{t_n}} \tag{S42}$$

$$= \sum_{\{n \,|\, t_n \geq t\}} \sum_k \kappa^{t_n - t} W_{kj}^{\pi,\mathrm{out}} \pi_k^{t_n} \left(\log \pi_k^{t_n} + H^{t_n}\right) . \tag{S43}$$

And thus, using a similar derivation as in equations (S15) to (S17), we find the estimate of the gradient of the loss function $E_H$ as:

$$\widehat{\frac{dE_H}{dW_{ji}}} = \sum_t \frac{\partial E_H}{\partial z_j^t} e_{ji}^t = \sum_{n,k} W_{kj}^{\pi,\mathrm{out}} \pi_k^{t_n} \left(\log \pi_k^{t_n} + H^{t_n}\right) \bar{e}_{ji}^{t_n} . \tag{S44}$$

Hence this adds a term of the form $L_j^{H,t}\ \bar{e}_{ji}^t$ to the final learning rule of *reward-based e-prop* exhibited in equations (36) and (37) in Methods such that it becomes:

$$\Delta W_{ji}^{\mathrm{rec}} = -\eta \sum_t \delta^t \mathcal{F}_\gamma \left( L_j^t\ \bar{e}_{ji}^t \right) - \eta L_j^{H,t}\ \bar{e}_{ji}^t \quad \text{with} \tag{S45}$$

$$L_j^{H,t} = c_H \sum_k B_{jk}^\pi \pi_k^t \left(\log \pi_k^t + H^t\right) . \tag{S46}$$

**Plasticity rule for the output weights and biases:** The gradient of $E$ with respect to the output weights can be computed directly from equation (32) without the theory of *e-prop*. However, it also needs to account for the sum over future rewards that is present in the term $R^t - V^t$. Using a similar derivation as in equations (S33)-(S35) the plasticity rule for these weights becomes:

$$\Delta W_{kj}^{\pi,\mathrm{out}} = -\eta \sum_t \left[ \delta^t \mathcal{F}_\gamma \Big( \left(\pi_k^t - \mathbb{1}_{a^t = k}\right) \mathcal{F}_\kappa(z_j^t) \Big) + \pi_k^t \left(\log \pi_k^t + H^t\right) \mathcal{F}_\kappa(z_j^t) \right] \tag{S47}$$

$$\Delta W_j^{\mathrm{V,out}} = \eta c_V \sum_t \delta^t \mathcal{F}_\gamma \left(\mathcal{F}_\kappa(z_j^t)\right) . \tag{S48}$$

Similarly, we find the update rules for the biases of the output neurons. We obtain for these: $\Delta b_k^{\pi,\mathrm{out}} = -\eta \sum_t \left[ \delta^t \mathcal{F}_\gamma \Big( \left(\pi_k^t - \mathbb{1}_{a^t=k}\right) \Big) + \pi_k^t \left(\log \pi_k^t + H^t\right) \right]$, and $\Delta b^{\mathrm{V,out}} = \eta c_V \sum_t \delta^t$.

#### S5.2 Simulation details: RL version of the task where temporal credit assignment is difficult (Fig. S5)

The task considered in this experiment was the same as in section S3.4, but while the task was there formulated as a supervised learning, the network is trained here using a reinforcement learning setup. A reward of 1 was given at the end of the trial, if the agent selected the side which had more cues than the other side, otherwise no reward was given. The network model remained the same as in the supervised setup. The result was computed over 10 different runs and is shown in Fig. S5: The task can be learnt by *reward-based e-prop.* Note that each run was characterized by a different random seed, similar to section S3.4.

**Details of the decision process:** In the reinforcement learning setup of the task, one binary action formalizes the decision of the agent ("left" of "right") at the end of the trial. This decision was sampled according to probabilities $\pi_k$ that were computed from the network output using a softmax operation (see section S5.1).

**Details of the learning procedure:** For learning, we simulated batches of 64 trials, and applied weight changes at the end of each batch. Independent of the learning method, we used Adam to implement the weight update, using gradients that were accumulated in 64 trials using a learning rate of $5 \cdot 10^{-3}$ and default hyperparameters [2]. For *random e-prop*, we sampled broadcast weights $B_{jk}$ from a Gaussian distribution with a mean of 0 and a variance of 1. To avoid an excessively high firing rate, regularization, as described in section S2.2, was applied with $c_{\text{reg}} = 0.1$ and a target firing rate of $f^{\text{target}} = 10$ Hz.

##### S5.3 Simulation details: Atari task (Fig. 4, Fig. 5)

**Details of the Atari simulator and of the input scheme:** For all our experiments we used the Arcade Learning Environment [17] to simulate various Atari games. This simulator takes the actions from the agent as input and advances the dynamics of the game by repeating the same action for 4 times. The simulator then produces a reward feedback along with the new video frame. The shape of the produced video frame is 160 pixels in width and 210 pixels in height and each pixel has 3 values that describes its color. We first converted the colored video frame into a gray-scale video frame and scaled it to a size of 84 by 84. The gray-scale values (assuming values between 0 and 255) were divided by 255. In our experiments, these observations were each given for 5 ms as input $\boldsymbol{x}^t, \ldots, \boldsymbol{x}^{t+4}$ to the spiking agent.

**Details of evaluation:** The results reported in Fig. 4 and 5 were computed over 5 different runs, where each run was characterized by a different random seed. For each run this resulted in different random initial weights of the agent and different random numbers for sampling the stochastic action. We chose to compare our spiking agents to the results reported in Table S3 of [18] because of the similarity of the learning approach. Note however that there exist some differences in the evaluation protocol:

1. the results reported in [18] are given as the 5 best runs out of 50, whereas we report the average over 5 runs;
2. their evaluation protocol involves human starting conditions, meaning that evaluation episodes are initialized with human game play, whereas we start episodes randomly with up to 30 No-op actions.

**Details of the network model:** The network architecture of the agent consisted of a spiking CNN followed by an LSNN. The spiking CNN consisted of 5512 LIF neurons for Pong (10928 LIF neurons for Fishing Derby respectively). These neurons were organized in two layers. Neurons in the first layer received input current resulting from a convolution of the video frame. The spikes of these neurons were propagated to a second layer of neurons using synapses with a convolutional weight structure. Biases were channel-specific for both convolutions, see Table S1 for all corresponding hyperparameters. All neurons in the two convolutional layers had a membrane time constant of $\tau_{m,\text{CNN}}$ ms and a threshold of $v_{\text{th,CNN}}$, see Table S1. We were forced to use a very high rate of input frames in order to keep the compute time of the learning experiments within the range of our computing resources. Therefore a new image was presented to the network every 5 ms. This forced us to set the membrane constants of the spiking neurons in the preprocessing CNN to the unbiologically low value of 1 ms. We conjecture that a simulation of the same learning processes with larger computing resources, where the presentation time of input frames can be multiplied with some factor F, say F = 20, the spiking CNN would be able to process these visual inputs with neurons whose membrane time constants are multiplied with a corresponding factor F, thereby bringing them into a biological range.

The LSNN consisted of 240 LIF neurons and 160 ALIF neurons for Pong (180 LIF neurons and 120 ALIF neurons for Fishing Derby respectively) and received spiking input from all neurons of the second convolutional layer. Membrane time constants $\tau_m$, baseline thresholds $v_{\text{th}}$, adaptation time constants $\tau_a$ and adaptation strengths $\beta$ are reported in Table S1. The refractory period was set to 5 ms. All synaptic delays in the LSNN were 1 ms. The output $\boldsymbol{y}^t, V^t$ was defined as the membrane voltage of output neurons, whose membrane time constants $\tau_{out}$ are reported in Table S1. These output neurons received spikes from all neurons in the network, including neurons in the convolutional layers.

**Details of the action set and of the action generation:** The set of valid actions was selected according to the game at hand. For Pong, we used a set of 3 valid actions: Up, Down and Stay (No op). For Fishing Derby, we used 9 possible action choices: Stay (No op), Fire, Up, Right, Left, Down, Up-right, Up-Left, Down-right and Down-left. Actions were sampled according to a discrete probability distribution, where the $k$-th action was sampled with a probability of $\pi_k^t = \text{softmax}_k(y_1^{t_n}, \ldots, y_K^t)$. Since the agent received the same video frame for 5 ms, actions were also generated only after every 5 ms.

**Details of the learning procedure:** The agent was trained consuming a total of $250 \cdot 10^6$ action generations, corresponding to $10^9$ video frames of the Atari simulator (actions are repeated 4 times internally). We computed gradients resulting from *reward-based symmetric e-prop* according to equation (S45). A common learning scheme is to use actors living in parallel copies of the environment that can regularly communicate gradients with each other during ongoing episodes [18]. But this scheme can hardly provide a model for learning in the brain. For this reason, we designed a schedule of episodes of increasing length and performed weight updates only after 32 completed episodes (64 for Fishing Derby, respectively). A single agent can implement this algorithm through local synaptic plasticity. However this scheme is equivalent to simulating these 32 independent episodes in parallel -but without communication between learning agents- instead of sequentially. This observation can be used to save simulation time.

For Pong the weights of the CNN are only trained to minimize the regularization of the neural activity, hence it does not require learning signals. For Fishing Derby, as every neuron is connected to the output neurons, we define broadcast weights $B_{jk}^{\pi,\text{CNN}}, B_{jk}^{V,\text{CNN}}$ for neurons in the CNN to be symmetric to the output weights: $B_{jk}^{\pi,\text{CNN}} = W_{kj}^{\pi,\text{out},\text{CNN}}, B_j^{V,CNN} = W_j^{V,\text{out},\text{CNN}}$.

Due to simulation constraints, we reset the fading memory filter $F_\gamma(L_j^t \bar{e}_{ji}^t)$ as well as the eligibility vectors $\boldsymbol{\epsilon}_{ji}^t$ to zero after every 500 ms for Pong (150 ms for Fishing Derby respectively). The weight updates were applied using Adam [2] with a learning rate of $10^{-3}$ (The parameter $\epsilon$ was set to 1). In Fishing Derby we clipped these gradients if exceeding a norm of 1000. The maximum episode length $T_{\max}$ was restricted to 1000 ms at the start of training, and was increased after every 8750 s by $\Delta T$, see Table S1. We used a discount factor of $\gamma = 0.998$. Note that an action is generated only after every 5 ms. The coefficient $c_V$ that trades off between $E_\pi$ and $E_V$ can be found in Table S1. We also applied entropy regularization according to equation (S44) to prevent premature convergence to suboptimal policies. The value of the coefficient $c_H$ that adds this loss $E_H$ to the objective is provided in Table S1. To avoid an excessively high firing rate, regularization was applied as described in S2.2. The target firing rate was given by $f^{\text{target}} = 20$ Hz. We used further regularization to prevent membrane voltages from assuming values for which $\psi_j$ is zero.

### S6 Evaluation of four variations of *e-prop* (Fig. S7)

We evaluate here the performance of four variations of *random e-prop*. In these variations, we used

- truncated eligibility traces for LIF neurons,
- global broadcast weights,
- temporally local broadcast weights, and
- a replacement of the eligibility trace by the corresponding term of the Clopath rule,

respectively. The considered task, whose implementation details are described in section S6.5, is an extension of the task used in [1]. In this task, an RSNN was trained to autonomously generate a 3 dimensional target signal for 1 second. Each dimension of the target signal was given by the sum of four sinusoids with random phases and amplitudes. Similar to [1], the network received a clock input that indicated the current phase of the pattern.

In Fig. S7a, we show the spiking activity of a randomly chosen subset of 20 out of the 600 neurons in the RSNN along with the output of the three output neurons after application of *random e-prop* for 1, 100, and 500 seconds, respectively. In this representative example, the network achieved a very good fit to the target signal (normalized mean squared error 0.01).

##### S6.1 A truncated eligibility trace for LIF neurons

A replacement of the term $\bar{z}_i^t$ with $z_i^t$ in equation (23) yields a performance that is reported in panel b of Fig. S7 as "Trunc. e-trace". Its performance is for the considered task only slightly worse than that of *random e-prop.*

##### S6.2 Global broadcast weights

Since 3-factor rules have primarily been studied so far with a global 3rd factor, we asked how the performance of *e-prop* would change if the same broadcast weight would be used for broadcast connections between all output neurons $k$ and network neurons $j$. We set this global broadcast weight equal to $\frac{1}{\sqrt{n}}$ with $n$ being the number of neurons in the network. Fig. 2 and Fig. S2 show together that the performance for the considered task is much worse than that of *random e-prop.* We have also tested this on TIMIT with LSNNs and found there an increase of the frame-wise error rate from 36.9% to 52% when replacing the broadcast weights of *random e-prop* with a global one. On the harder version of same task, the error rate at the sequence level increased from 34.7% to 60%.

##### S6.3 Temporally local broadcast weights

One can train RNNs also by applying the broadcast alignment method of [19] and [20] for feed-forward networks to the unrolled version (see Fig. 1b) of the RNN. In contrast to *e-prop*, this approach suggests to draw new random broadcast weights for each layer of the unrolled network, i.e., for each time step of the RNN. Fig. S7c shows that this variation of *random e-prop* performs much worse. However an intermediate version where the random broadcast weights are redrawn every 20 ms performs about equally well as *random e-prop* for the considered task.

##### S6.4 Replacing the eligibility trace by the corresponding term of the Clopath rule

The dependence of the synaptic plasticity rules from *e-prop* on the postsynaptic membrane potential through the pseudo-derivative in the eligibility traces yields some similarity to some previously proposed rules for synaptic plasticity, such as that of [21], which were motivated by experimental data on the dependence of synaptic plasticity on the postsynaptic membrane potential. We therefore tested the performance of *random e-prop*, where the eligibility trace was replaced by the corresponding term from the "Clopath rule":

$$[v_j^t - v_{\mathrm{th}}^+]^+ [\bar{v}_j^t - v_{\mathrm{th}}^-]^+ \bar{z}_i^{t-1} \ , \tag{S49}$$

where $\bar{v}_j^t$ is an exponential trace of the post synaptic membrane potential, with a time constant of 10 ms chosen to match their data. $[\cdot]^+$ is the rectified linear function. The thresholds $v_{\mathrm{th}}^-$ and $v_{\mathrm{th}}^+$ were $\frac{v_{\mathrm{th}}}{4}$ and 0 respectively. Fig. S7b shows that the resulting synaptic plasticity rule performed quite well.

##### S6.5 Simulation details: pattern generation task

The performance in this task is reported as a normalized mean squared error (nmse) that we defined for this task as: nmse $= \frac{\sum_{t,k}(y_k^t - y_k^{*,t})^2}{\sum_{t,k}(y_k^{*,t} - \bar{y}_k^*)^2}$, where we set $\bar{y}_k^* = \frac{1}{T}\sum_t y_k^{*,t}$.

**Details of the network model and of the input scheme:** We used a network that consisted of 600 LIF neurons. Each neuron had a membrane time constant of $\tau_m = 20$ ms and a refractory period of 3 ms. The firing threshold was set to $v_{\text{th}} = 0.41$. Output neurons used a membrane time constant of $\tau_{out} = 20$ ms. The network received input from 20 input neurons, divided into 5 groups, which indicated the current phase of the target sequence similar to [1]. Neurons in group $i \in \{0, 4\}$ produced 100 Hz regular spike trains during the time interval $[200 \cdot i, 200 \cdot i + 200)$ ms and were silent at other times.

**Details of the target pattern:** The target signal had a duration of 1000 ms and each component was given by the sum of four sinusoids, with fixed frequencies of 1 Hz, 2 Hz, 3 Hz, and 5 Hz. At the start of learning, the amplitude and phase of each sinusoid in each component was drawn uniformly in the range $[0.5, 2]$ and $[0, 2\pi]$ respectively. This signal was not changed afterwards.

**Details of the learning procedure:** For learning, we computed gradients after every 1 second of simulation, and carried out the weight update using Adam [2] with a learning rate of $3 \cdot 10^{-3}$ and default hyperparameters. After every 100 iterations, the learning rate was decayed by a factor of 0.7. For *random e-prop*, the broadcast weights $B_{jk}$ were sampled from a Gaussian distribution with a mean of 0 and a variance of $\frac{1}{n}$, where $n$ is the number of network neurons.

Firing rate regularization, as described in section S2.2, was applied with $c_{\text{reg}} = 0.5$ and a target firing rate of $f^{\text{target}} = 10$ Hz.

| | Pong | Fishing Derby |
|---|---|---|
| $\tau_m$ | 15 ms | 10 ms |
| $v_{th}$ | 1 | 1 |
| $\tau_a$ | 70 ms | 70 ms |
| $\beta$ | 0.1 | 0.1 |
| $\tau_{out}$ | 2 ms | 4 ms |
| $v_{\mathrm{th,CNN}}$ | 0.1 | 0.1 |
| $\tau_{m,\mathrm{CNN}}$ | 1 ms | 1 ms |
| $c_V$ | 1 | 0.5 |
| $c_H$ | 0.0025 | 0.006 |
| $c_{\mathrm{reg}}$ | 50 | 10 |
| $c_{\mathrm{reg},V}$ | $10^{-4}$ | $10^{-3}$ |
| $\Delta T$ | $\left(\left(\sqrt[3]{\frac{T_{\max}}{1\ s}\cdot 1000}+1\right)^3-\frac{T_{\max}}{1\ s}\cdot 1000\right)$ ms | 1000 ms |
| First convolutional Layer | | |
| Kernel structure | $8\times 8\times 1\times 8$ | $8\times 8\times 1\times 16$ |
| Stride | 4 | 4 |
| Padding | VALID | SAME |
| Number of neurons | 3200 | 7056 |
| Second convolutional Layer | | |
| Kernel structure | $4\times 4\times 8\times 8$ | $4\times 4\times 16\times 32$ |
| Stride | 1 | 2 |
| Padding | VALID | SAME |
| Number of neurons | 2312 | 3872 |

Table S1: Table of hyperparameters used in Atari tasks

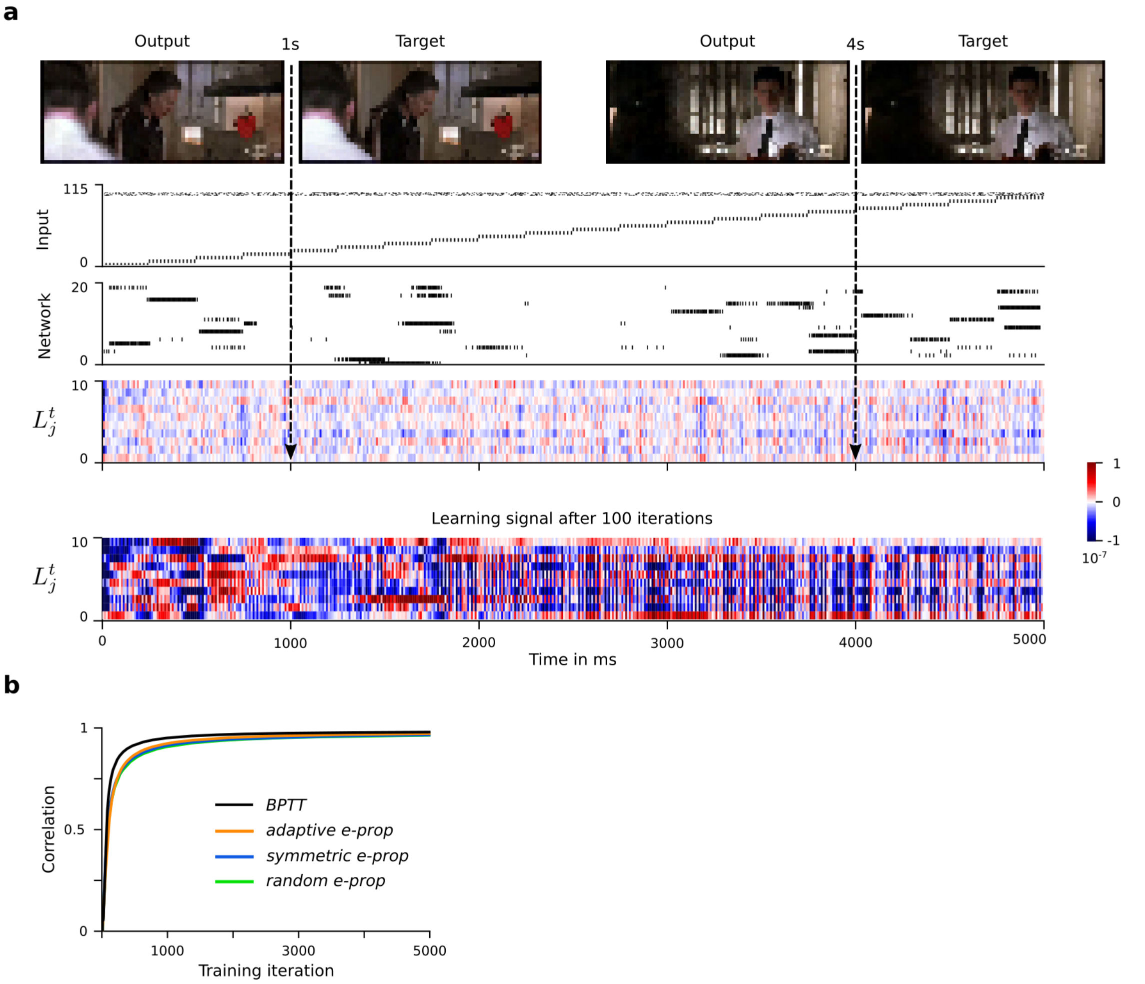

Figure S1: **Performance comparison of *BPTT* and *e-prop* on an extension of the episodic memory task from** [1]**.** The LSNN had to generate at each time step the values of all pixels that described the video frame of the movie at that time step, see section S3.3. **a**) Input spikes, network activity (for 20 sample neurons), learning signals, and network outputs (at 1s and 4s, shown at the top) of an LSNN after 1000 training iterations. For comparison we also show learning signals after just 100 iterations, where their amplitude is still large. **b**) Performance of *BPTT* and *e-prop*.

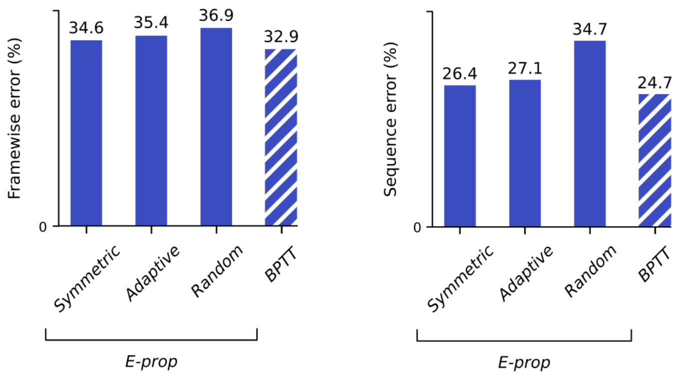

Figure S2: **Comparison of learning algorithms for training LSNNs on the TIMIT task.** Performance of *BPTT* and the three versions of *e-prop* on frame-wise phoneme classification (left) and for phoneme sequence recognition (right).

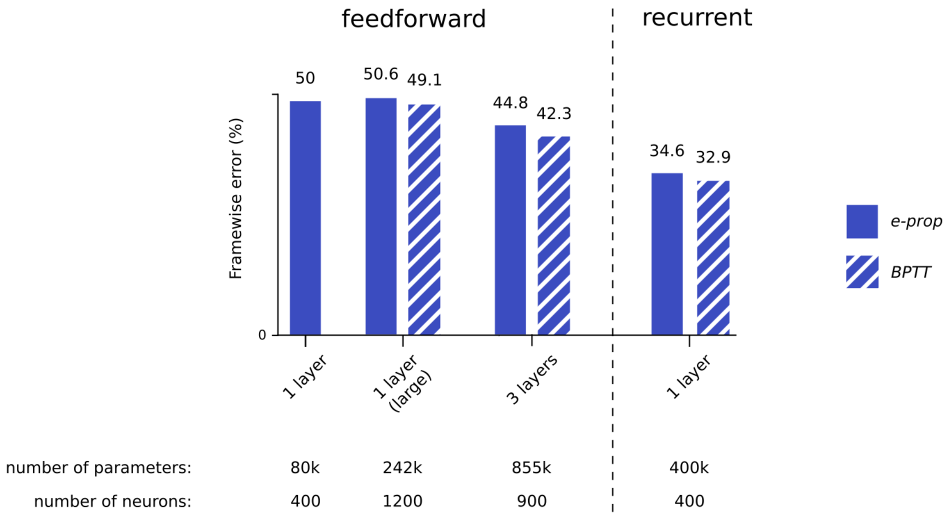

Figure S3: **Performance of *e-prop* on the framewise TIMIT task for variations of the LSNN from Fig. 2 without recurrent connections (left of dashed line).** The result of the left-most bar was achieved with the same hyperparameters as used in Figure 2. For the other feedforward architectures we modified hyperparameters to optimize the performance. The results of Fig. 2 **c)** for LSNNs with recurrent connections are redrawn on the right of the dashed line. One sees that recurrent connections are essential for achieving good performance on this task.

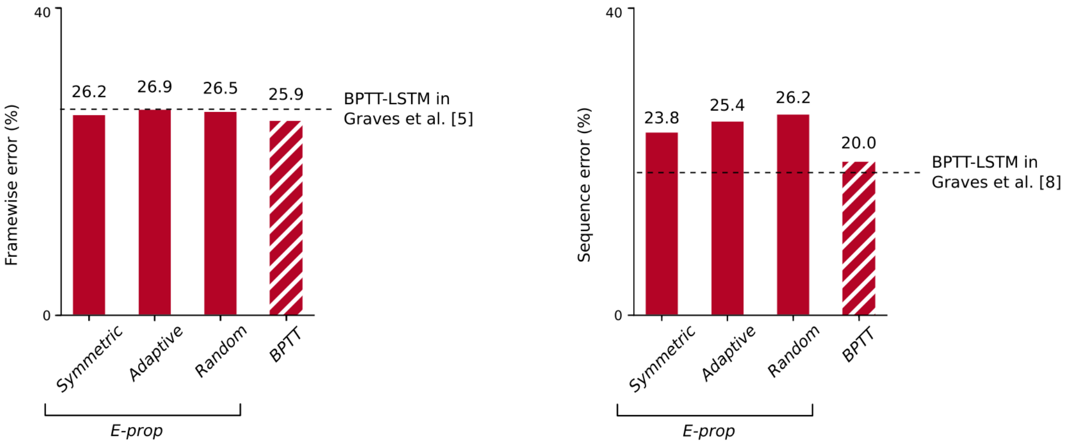

Figure S4: **LSTM networks trained with *BPTT* and *e-prop* on the TIMIT task.** Performance of *BPTT* and the three versions of *e-prop* on frame-wise phoneme classification (left) and for phoneme sequence recognition (right). One sees that *e-prop* approximates *BPTT* performance also for non-spiking neural networks. The *BPTT* baselines aim at reproducing the results obtained with LSTM networks in [6] and [9]. The network architectures and audio pre-processing settings are taken from [6, 7] for framewise phoneme classifcation and from [9] for phonemene sequence transcription. In comparison to the *BPTT*-LSTM baselines that we could achieve in this way, 26.9% framewise error rate was reported in [6] and 18.6% sequence error rate was reported in [9].

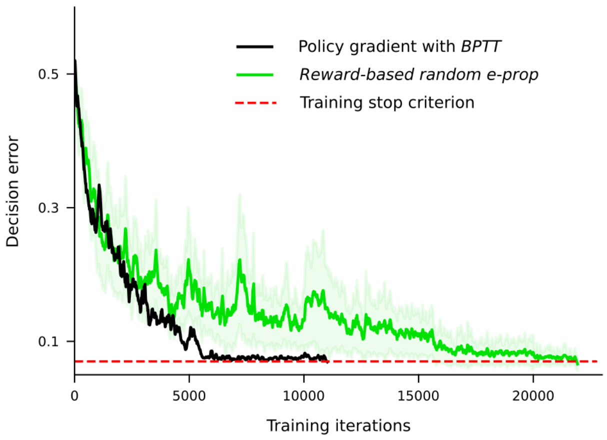

Figure S5: **Performance of *reward-based random e-prop* and *BPTT* for the RL version of the task from Fig. 3, applied to an LSNN consisting of 50 LIF and 50 ALIF neurons.**

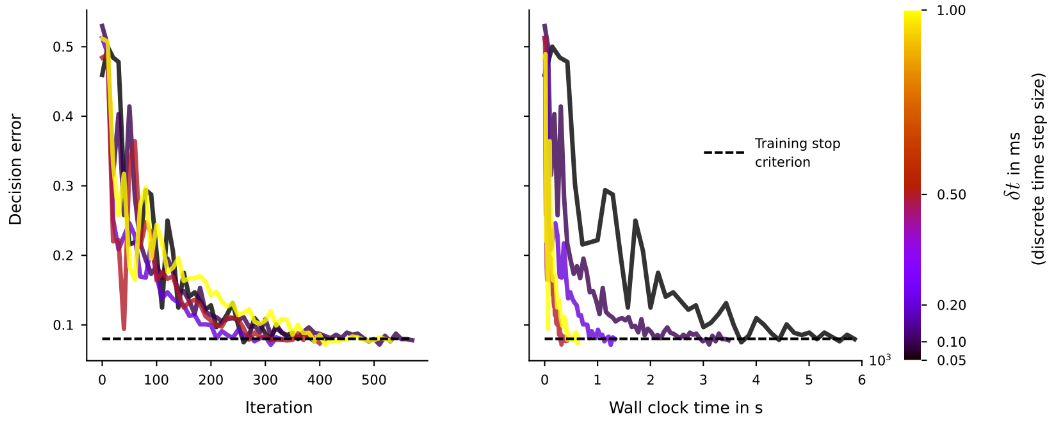

Figure S6: **Impact of the length of the simulation time step on the learning performance of *e-prop* for the task of Fig. 3 (simplified version with 5 cues instead of 7).** Decision error for this task is shown as function of the number of training iterations, and as function of wall clock time. The results are averaged over 5 different seeds. One sees that the length of the simulation time step has no visible impact on the learning performance of *e-prop*. However, smaller simulation time steps lead to substantially larger simulation time

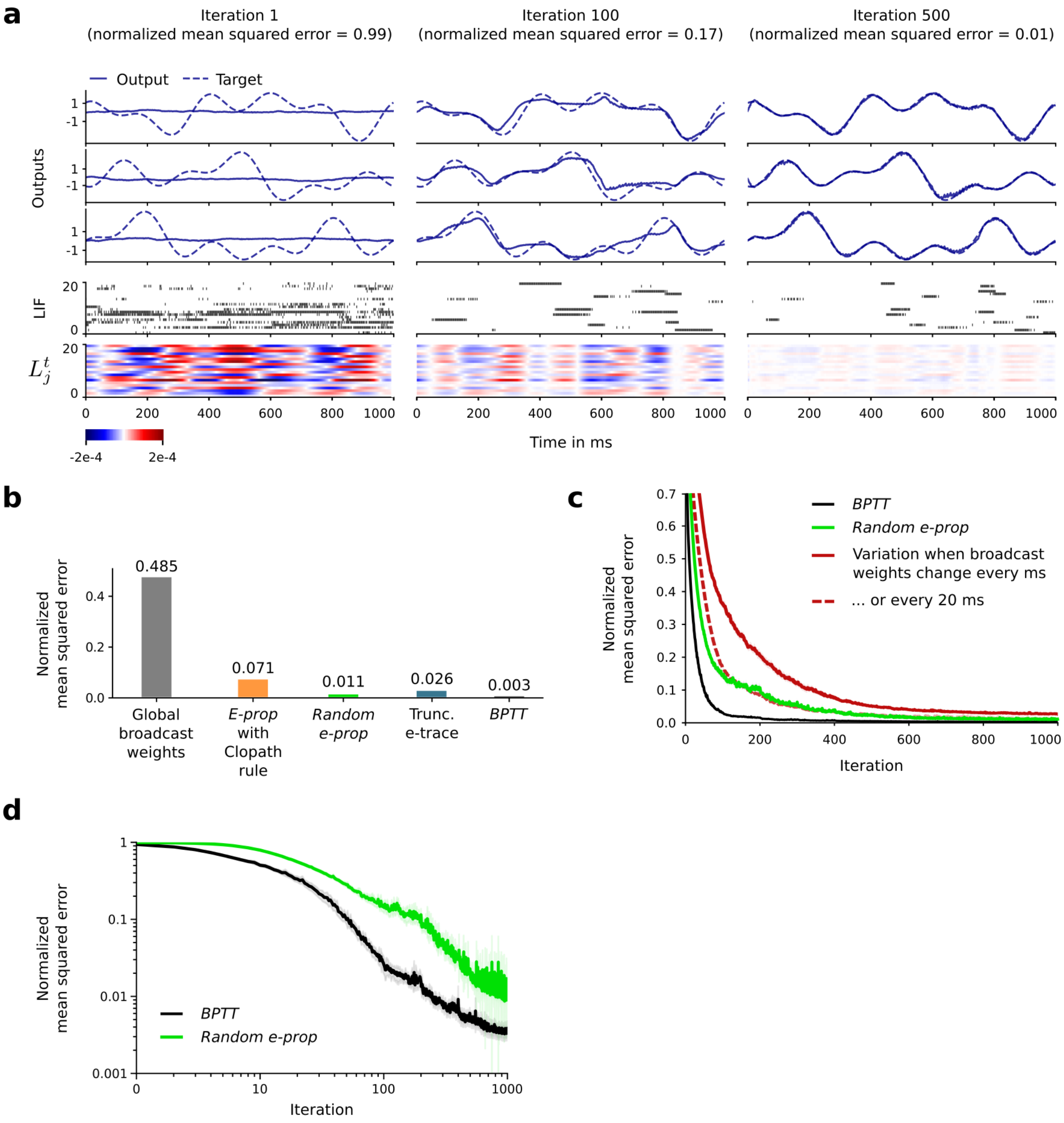

Figure S7: **Evaluation of several variants of *random e-prop*** **a**) The task is a classical benchmark task for learning in recurrent SNNs: learning to generate a target pattern, extended here to the challenge to simultaneously learn to generate 3 different patterns, which makes credit assignment for errors more difficult. Learning performance with *random e-prop* is shown after training for 1, 100, 500 s. **b**) Normalized mean squared error of several learning algorithms for this task after 500 s of training. "Clopath rule" denotes a replacement of the eligibility trace of *random e-prop* by a corresponding term proposed in [21] based on experimental data. **c**) Learning curves for variations of *random e-prop* with temporally local broadcast weights. Results were averaged over 20 runs. **d**) Comparison of learning curves for *BPTT* and random *e-prop* with logarithmic scale.

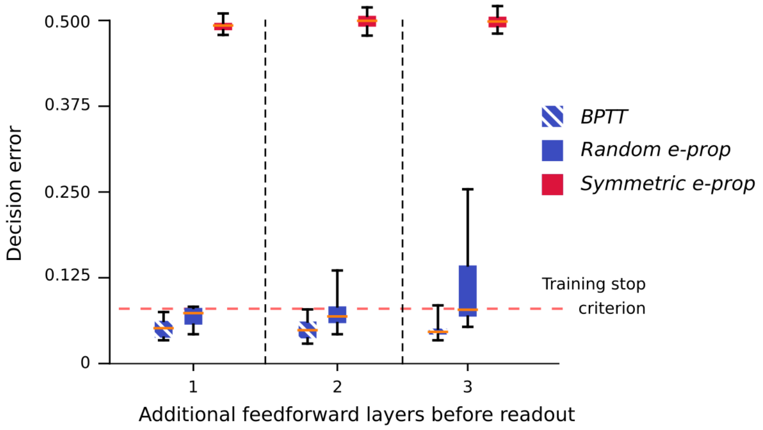

Figure S8: **Test of a prediction of our analysis when *e-prop* is likely to approximate the performance of *BPTT*, and when not.** We blocked in the learning task of Fig. 3 route i for gradient information by inserting 1 to 3 feedforward layers (completely connected from one layer to the next) between the RSNN and the readouts $y$ of Fig. 2a. All neurons in the RSNN were connected to the first feedforward layer, and all neurons in the last layer to the readout neurons. The decision error is reported for a distribution of runs with different initialization weights. The upper and lower whiskers are the minimum and the maximum, the orange bar is the median and the box extends from the lower to the upper quartile of the data. As predicted, the feedforward layers drastically reduce the performance of *symmetric e-prop*, but not of *BPTT*. Thus for successful training of a RSNN by *symmetric e-prop* it is essential that all or most neurons in the RSNN have a synaptic connection to readout neurons. *Random e-prop* works still quite well, but the high variance of its performance for 3 intermediate feedforward layers suggests some learning instability. See section S3.4 for details of this experiment.

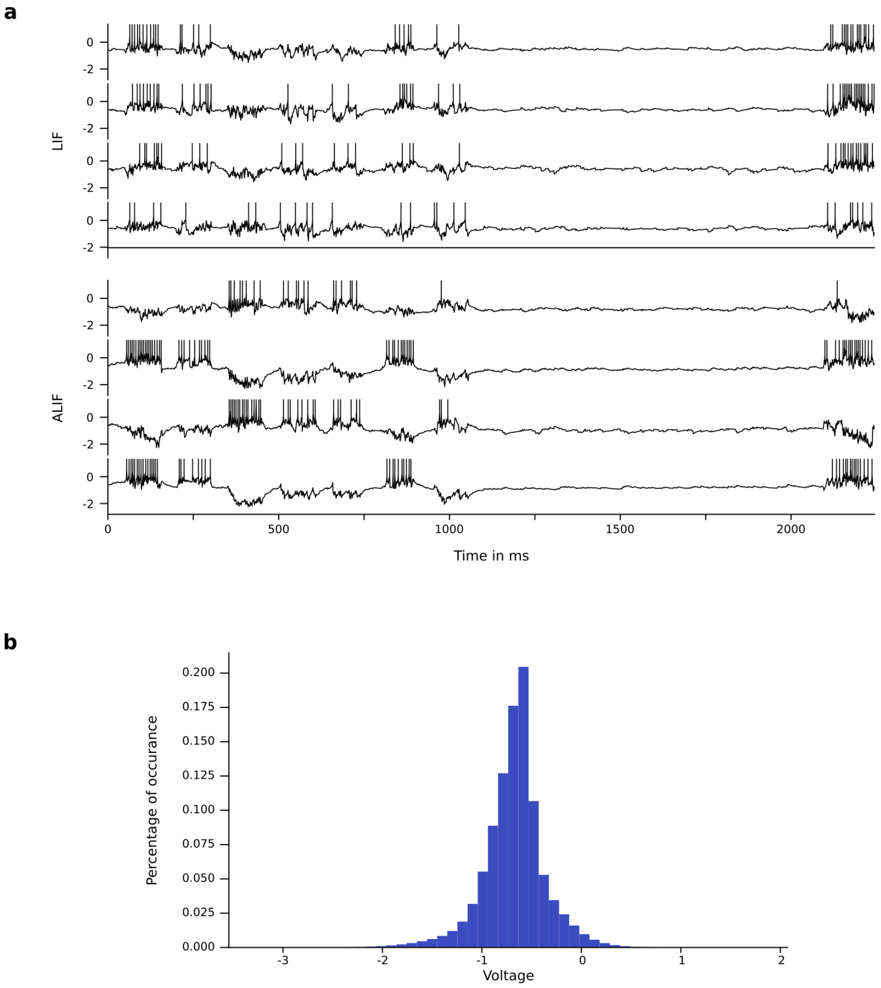

Figure S9: **Further details on membrane potentials of neurons in the network of Fig. 3** **a**. Traces of voltage – firing threshold $(v_j^t - A_j^t)$ for 4 randomly selected LIF and 4 ALIF neurons. **b**) Histogram of these values for all neurons in the network and all time points $t$.

**Movie S1: Task from Fig. 3 with difficult temporal credit assignment**

Rodent task from [14] and [22] that requires long-term credit assignment for learning: a rodent has to learn to run along a linear track in a virtual environment, where it encounters several cues on the left and the right side along the way. It then has to run through a corridor without cues (giving rise to delays of varying lengths). At the end of the corridor, the rodent has to turn to either the left or the right side of a T-junction, depending on which side exhibited more cues along the way.

**Movie S2: Dynamics of *BPTT* for the task from Fig. 3 with difficult temporal credit assignment**

Dynamics of *BPTT* for the task where temporal credit assignment is difficult: First, a simulation of the network has to be carried out in order to produce the network state of all neurons for all time steps. After that the loss function $E$ can be evaluated. Then the simulated network activity is replayed backwards in time to assign credit to particular spikes that occurred before the loss function became non-zero. One sees that the slow time constants that are present in the dynamics of adapting thresholds of adaptive LIF (ALIF) neurons result in slowly decaying non-vanishing gradients during the backpropagation through time ("highways into the past"). In contrast, for LIF neurons the backpropagated gradients vanish rather quickly.

**Movie S3: Dynamics of *e-prop* for the task from Fig. 3 with difficult temporal credit assignment**

The computation of the LSNN is accompanied by the computation of synapse specific eligibility traces. An error in the computation only becomes apparent during the so-called decision period at the end of a trial. In this last phase, a learning signal ($L_j$) that transmits deficiencies of the network output is provided separately to each neuron. As can be seen from the video, synapses that project to neurons with adapting thresholds (ALIF neurons) still have non-vanishing eligibility traces during the last phase ("highways into the future"), and hence can be combined with the learning signals at that time to implement long-term credit assignment.

**Movie S4: Solution of the episodic memory task from [1] by an LSNN trained with *e-prop***

Extension of episodic memory task from [1] trained with *random e-prop*. The top row presents the actual movie clip, and the output produced by the trained LSNN. The middle row shows the input that is presented to the network: a channel that indicates which of the three learned clips had to be replayed, and an array of input neurons that indicate the current timing in the clip. The bottom row shows the spiking activity of a subset of the neurons in the LSNN (20 neurons out of 1000). As can be seen, the network learned via *e-prop* to distinguish well between the different clips and also, the LSNN was able to deal with scene cuts, which require the network to change its output abruptly.

**Movie S5: Network dynamics, synaptic plasticity, and performance of an LSNN trained by *reward-based e-prop* to win the Atari game Pong**

A trial of the Atari Pong task after training the network with *reward-based symmetric e-prop*: The video frames of the game screen (right) are preprocessed using a spiking CNN and provided as input to the LSNN, resulting in spiking activity (bottom left: LIF, ALIF). Output neurons predict future rewards, and the probability of taking actions (Move up, Move down, Stay), shown top left. Learning dynamics are induced when reward prediction errors (green) are combined with a decaying product of local eligibility traces and action feedback (blue). This results in weight changes at synapses (red), shown middle left.
